# Validity and cultural generalisability of a 5-minute AI-based, computerised cognitive assessment in Mild Cognitive Impairment and Alzheimer’s Dementia

**DOI:** 10.1101/2021.04.01.437840

**Authors:** Hadi Modarres, Chris Kalafatis, Panos Apostolou, Haniye Marefat, Mahdiyeh Khanbagi, Hamed Karimi, Zahra Vahabi, Dag Aarsland, Seyed-Mahdi Khaligh-Razavi

## Abstract

**INTRODUCTION:** Early detection and monitoring of mild cognitive impairment (MCI) and Alzheimer’s Disease (AD) patients are key to tackling dementia and providing benefits to patients, caregivers, healthcare providers and society. We developed the Integrated Cognitive Assessment (ICA); a 5-minute, language independent computerised cognitive test that employs an Artificial Intelligence (AI) model to improve its accuracy in detecting cognitive impairment. In this study, we aimed to evaluate the generalisability of the ICA in detecting cognitive impairment in MCI and mild AD patients.

**METHODS:** We studied the ICA in a total of 230 participants. 95 healthy volunteers, 80 MCI, and 55 participants with mild AD completed the ICA, the Montreal Cognitive Assessment (MoCA) and Addenbrooke’s Cognitive Examination (ACE) cognitive tests.

**RESULTS:** The ICA demonstrated convergent validity with MoCA (Pearson r = 0.58, p<0.0001) and ACE (r = 0.62, p<0.0001). The ICA AI model was able to detect cognitive impairment with an AUC of 81% for MCI patients, and 88% for mild AD patients. The AI model demonstrated improved performance with increased training data and showed generalisability in performance from one population to another. The ICA correlation of 0.17 (p=0.01) with education years is considerably smaller than that of MoCA (r=0.34, p<0.0001) and ACE (r=0.41, p<0.0001) which displayed significant correlations. In a separate study the ICA demonstrated no significant practice effect observed over the duration of the study.

**DISCUSSION:** The ICA can support clinicians by aiding accurate diagnosis of MCI and AD and is appropriate for large-scale screening of cognitive impairment. The ICA is unbiased by differences in language, culture and education.

## 1. Introduction

Neurodegenerative disorders, including dementia and Alzheimer’s Disease (AD), continue to represent a major social, healthcare and economic burden, worldwide.^1^ AD is the most common type of dementia with mild cognitive impairment (MCI) being a pre-dementia condition with a prevalence ranging from 16% to 20% of the population in those between 60 and 89 years old.^2^ Of patients suffering from MCI, 5-15% progress to dementia every year.^3^ These diseases remain underdiagnosed or are diagnosed too late, potentially resulting in less favourable health outcomes as well as higher costs on healthcare and social care systems.^4^

In anticipation of disease-modifying treatments for MCI and AD,^5,6^ the importance of early diagnosis has become increasingly pressing.^7^ There is accumulating evidence that early detection provides cost savings for health care systems and is an achievable goal,^8,9^ and accurate patient selection for disease-modifying treatments is cost-effective and will improve clinical outcomes.^5^

Timely identification and diagnosis is considered to be key to tackling dementia offering multiple benefits to patients, families and caregivers, healthcare providers, as well as society as a whole.^10^ If achieved, early detection can aid the deceleration of the progression of the disease.^8^ Early disease identification can consolidate preventative efforts through modifiable lifestyle factors to limit the progression of the disease.^11^

The available neuroimaging and fluid biomarkers of neurodegeneration are not easily accessible or scalable as health services cannot provide them routinely.^5^ As a result, neuropsychological assessments remain the mainstay of dementia diagnosis. Current routinely used neuropsychological assessments are invariably paper-based, language and education-dependent, have ceiling effects and require substantial clinical time to administer.

They lack reliability in preclinical stages of dementia,^12,13^ and are prone to classification errors.^14^ Such tests are typically subject to practice effects,^15^ which may lead to incorrect estimates of age-related changes. Invariably, these tests require administration by a clinician and therefore are not appropriate for remote measurements or longitudinal monitoring.

Computerized cognitive tests constitute promising tools for the early detection of clinically relevant changes in MCI and AD sufferers.^7,16–18^ Computer and mobile device-based tests have been investigated to some extent, with several showing good diagnostic accuracy entering clinical use.^13,19,20^ Devices such as smartphones and tablets offer a nearly unlimited number of applications that can be combined for the comprehensive assessment of cognition and brain health in community-dwelling users.

Digital biomarkers can help identify and monitor subtle cognitive changes in individuals at risk of developing dementia.^21^ Furthermore such tests are able to benefit from advanced analytics for more accurate and personalised results. Artificial intelligence (AI) and machine learning are being increasingly used for applications in Dementia research.^7^ These tools have also been applied to detection of dementia using electronic health records and neuroimaging data.^22–24^ Recent evidence suggests that three-quarters of older people are open to new technology to make an accurate and early diagnosis and would agree to using computer or smartphone tasks that monitor day-to-day life.^25^

Computerised cognitive assessments have primarily been designed to mirror traditional pen-and-paper tests, ultimately missing the opportunity to obtain additional diagnostically useful cognitive information.^26^ Moreover, current computerised cognitive assessments have been struggling with facing trade-offs in terms of sensitivity and specificity, required effort and adherence, while others have been criticised for testing their technology with younger adults.^27^

Significantly, not much attention has been paid to personalised care in order to adapt to the individual’s cognitive and functional characteristics, such as offering tailored information to the needs of the patients.^28^

We developed the Integrated Cognitive Assessment (ICA) which is a 5-minute, self-administered, computerised cognitive assessment tool based on a rapid categorisation task that employs AI to detect cognitive impairment. The ICA is self-administered and independent of language.^29,30^ We aimed to evaluate the generalisability of the ICA in detecting cognitive impairment in MCI and mild AD patients. We recruited participants from two cohorts in different continents. We hypothesize that the AI-model employed for ICA can be generalised across demographically different patient populations.

To measure the convergent validity of the ICA with standard of care cognitive tests we compared the ICA with the Montreal Cognitive Assessment (MoCA) and Addenbrooke’s Cognitive Examination (ACE). We investigated the level of education bias between the cognitive assessments.

We also report the effects of repeated exposure to the test in healthy participants (learning bias).

## 2. Methodology

### 2.1. The ICA test description

The ICA test is a rapid visual categorization task with backward masking, and has been described in detail in previous publications.^29,30^ The test takes advantage of the human brain’s strong reaction to animal stimuli.^31–33^ One hundred natural images (50 of animals and 50 of not containing an animal) of various levels of difficulty are selected and are presented to the participant in rapid succession as shown in Figure S1 in the supporting information (SI).

Images are presented at the centre of the screen at 7° visual angle to the participant. In some images the head or body of the animal is clearly visible to the participants, which makes it easier to detect. In other images the animals are further away or otherwise presented in cluttered environments, making them more difficult to detect.

The strongest categorical division represented in the human higher level visual cortex appears to be that between animals and inanimate objects.^34,35^ Studies also show that on average it takes about 100 ms to 120 ms for the human brain to differentiate animate from inanimate stimuli.^32,36,37^ Following this rationale, each image is presented for 100 ms followed by a 20 millisecond inter-stimulus interval (ISI), followed by a dynamic noise mask (for 250 ms), followed by subject’s categorization into animal vs non-animal. Shorter periods of ISI can make the animal detection task more difficult and longer periods reduce the potential use for testing purposes as it may not allow for the detection of less severe cognitive impairments. The dynamic mask is used to remove (or at least reduce) the effect of recurrent processes in the brain.^38,39^ This makes the task more challenging by reducing the ongoing recurrent neural activity that could artificially boost the subject’s performance; it further reduces the chances of learning the stimuli. For more information about rapid visual categorization tasks refer to Mirzaei et al., (2013).^40^

Grayscale images are used to remove the possibility of color blindness affecting participants’ results. Furthermore, color images can facilitate animal detection solely based on color,^41,42^ without fully processing the shape of the stimulus. This could have made the task easier and less suitable for detecting mild cognitive deficits.

The ICA test begins with a different set of 10 trial images (5 animal, 5 non-animal) to familiarize participants with the task. If participants perform above chance (>50%) on these 10 images, they will continue to the main task. If they perform at chance level (or below), the test instructions are presented again, and a new set of 10 introductory images will follow. If they perform above chance in this second attempt, they will progress to the main task. If they perform below chance for the second time the test is restarted.

#### Backward masking

To construct the dynamic mask, following the procedure in,^43^ a white noise image was filtered at four different spatial scales, and the resulting images were thresholded to generate high contrast binary patterns. For each spatial scale, four new images were generated by rotating and mirroring the original image. This leaves us with a pool of 16 images. The noise mask used in the ICA test was a sequence of 8 images, chosen randomly from the pool, with each of the spatial scales to appear twice in the dynamic mask.

The ICA primarily tests Information processing speed (IPS) and engages higher-level visual areas in the brain for semantic processing, i.e. distinguishing animal vs. non-animal images,^29^ which is the strongest categorical division represented in the human higher-level visual cortex.^44^ IPS underlies many areas of cognitive dysfunction^45,46^ and is one of the key subtle, early changes that is slowed down in pre-symptomatic Alzheimer’s disease.^47^ This is because the speed with which an individual performs a cognitive task is not an isolated function of the processes required in that task, but also a reflection of their ability to rapidly carry out many different types of processing operations.

In the case of the ICA, these operations include transferring visual information through the retina to higher level visual areas i.e. sensory speed, processing the image representation in the visual system to categorize it into animal or non-animal (i.e. cognitive speed), and then translating this into a motor response i.e. motor speed.

### 2.2. Reference pen-and-paper cognitive tests

#### Montreal Cognitive Assessment

MoCA is a widely used screening tool for detecting cognitive impairment, typically in older adults.^48^ The MoCA test is a one-page 30-point test administered in approximately 10 minutes.

#### Addenbrooke’s Cognitive Examination (ACE)

The ACE was originally developed at Cambridge Memory Clinic.^49,50^ ACE assesses five cognitive domains: attention, memory, verbal fluency, language and visuospatial abilities. On average, the test takes about 20 minutes to administer and score.

ACE-R is a revised version of ACE that includes MMSE score as one of its sub-scores.^51^ ACE-III replaces elements shared with MMSE and has similar levels of sensitivity and specificity to ACE-R. ^52^

### 2.3. Study design

We aimed at studying the ICA across a broader spectrum of geographical locations with differences in language and culture to test the generalisability of the ICA. For analytical purposes we combined participants from two cohorts in order to study the ICA in one demographically diverse population.

See Table 1 for a summary of the demographic characteristics of recruited participants.

**Table 1.** Summary of demographic information, and cognitive test scores of recruited participants

|  |  |  |  | Age |  | Education<br>years |  | MoCA |  | ACE |  | ICA Index |  |
| --- | --- | --- | --- | --- | --- | --- | --- | --- | --- | --- | --- | --- | --- |
| Cohort | Diagnosis | Count | Female<br>(%) | Mean | SD | Mean | SD | Mean | SD | Mean | SD | Mean | SD |
| Cohort 1 | Healthy | 33 | 57.6 | 63.6 | 6.7 | 14.2 | 4.7 | 25.9 | 3.0 | 92.1 | 6.7 | 66.1 | 7.7 |
|  | MCI | 27 | 55.6 | 66.0 | 7.0 | 14.2 | 5.7 | 23.7 | 2.9 | 87.3 | 7.3 | 57.8 | 8.1 |
|  | mild AD | 13 | 46.2 | 69.8 | 9.4 | 11.2 | 4.0 | 16.6 | 5.8 | 68.5 | 14.3 | 41.6 | 14.4 |
| Cohort 2 | Healthy | 62 | 54.8 | 68.5 | 7.6 | 14.3 | 4.2 | 28.3 | 1.8 | 95.7 | 3.1 | 63.7 | 8.7 |
|  | MCI | 53 | 43.4 | 71.5 | 7.9 | 12.5 | 2.7 | 23.5 | 2.9 | 84.1 | 6.9 | 54.7 | 11.9 |
|  | mild AD | 42 | 50.0 | 71.6 | 7.4 | 13.3 | 3.2 | 20.2 | 3.0 | 76.1 | 8.1 | 46.9 | 15.5 |
| Combined | Healthy | 95 | 55.8 | 66.8 | 7.6 | 14.3 | 4.4 | 27.5 | 2.6 | 94.4 | 4.9 | 64.5 | 8.4 |
|  | MCI | 80 | 47.5 | 69.6 | 8.0 | 13.1 | 4.0 | 23.6 | 2.9 | 85.2 | 7.2 | 55.7 | 10.8 |
|  | mild AD | 55 | 49.1 | 71.2 | 7.9 | 12.8 | 3.5 | 19.3 | 4.1 | 74.3 | 10.3 | 45.6 | 15.3 |

#### 2.3.1. Cohort 1

73 (33 Healthy, 27 MCI, 13 mild AD) participants completed the ICA test, MoCA and ACE-R in the first assessment. The participants were non-English speakers, with instructions for the cognitive assessments provided in Farsi and were recruited from Neurology outpatients at the Royan Research Institute, Iran during a single visit. All assessment scales were carried out by a trained Psychologist or a study doctor.

All diagnoses were made by a consultant neurologist (Z.V) according to diagnostic criteria described by the working group formed by the National Institute of Neurological and Communicative Disorders and Stroke (NINCDS) and the Alzheimer’s Disease and Related Disorders Association (ADRDA) and the National Institute on Aging and Alzheimer’s Association (NIA-AA) diagnostic guidelines.^53^ All study participants had previously had an MRI-head scan, blood tests and physical examination as part of the diagnostic procedure.

The study was conducted at Royan institute. The study was conducted according to the Declaration of Helsinki and approved by the local ethics committee at Royan Institute. The inclusion exclusion criteria are listed in the Supplementary Information (SI).

#### 2.3.2. Cohort 2

157 (62 Healthy, 53 MCI, 42 mild AD) participants with a clinical diagnosis completed the ICA test, MoCA and ACE-III. The ICA test was taken on an iPad. Participants of age-range 55-90 were included in this study. The study was conducted at 6 NHS sites in the UK in participants recruited from NHS memory clinics. All assessment scales were administered by trained Psychologists or Nurses. Ethics approval was received from the London Dulwich Research Ethics Committee.

All diagnoses were made by a memory clinic consultant psychiatrist according to the same diagnostic criteria as in Cohort 1. The diagnostic procedure included an MRI-head scan, blood tests and physical examination for all participants. The eligibility criteria are listed in the SI. One additional inclusion criterion for Cohort 2 required an ACE-III score of >=90 for healthy participants. Cognitive assessments were performed either in the clinic, or via at home visits. Approximately 51% of assessments were conducted via home visits and 49% in the clinic in a single visit.

Inclusion criteria were common for both cohorts and refer to individuals with normal or corrected-to-normal vision, without severe upper limb arthropathy or motor problems that could prevent them from completing the tests independently (see SI). For each participant, information about age, education and gender was also collected. Informed written consent was obtained from all participants.

Spectrum bias, whereby the subjects included in the study do not include the complete spectrum of patient characteristics in the intended use population^54^ has been avoided in Cohort 1 and Cohort 2 by recruiting participants according to a sampling matrix and at the mild stage of Alzheimer’s Dementia. Therefore, the ICA performance metrics reported in this study are relative to detecting cognitive impairment in a population with less severe impairment.

### 2.4. Analysis methods

#### 2.4.1. Accuracy, speed and summary ICA Index calculation

The raw data from the ICA is composed of reaction time and categorisation accuracy on the images. This data was used to calculate summary features such as overall accuracy, and speed using the same methodology as described previously.^29,30^

Accuracy is defined as follows:

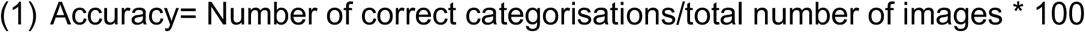

Speed is defined based on participant’s response reaction times in trials they responded correctly:

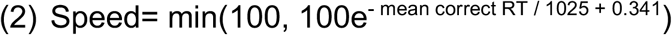

A summary ICA Index, is calculated as follows:

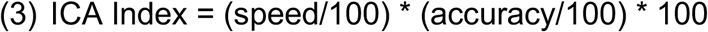

The ICA Index describes the raw test result, incorporating speed and accuracy, the two main elements of the ICA test.

#### 2.4.2. ICA AI model

The AI model utilizes inputs from accuracy and speed of responses to the ICA rapid categorisation task (with the ICA Index as an input feature), as well as age, and outputs an indication of likelihood of impairment (AI probability) by comparing the test performance and age of a patient to those previously taken by healthy and cognitively impaired individuals. The AI model is able to achieve an improved classification accuracy relative to using any single feature from the ICA test.

A probability threshold value of 0.5 was used to convert the AI probability to the AI prediction of healthy or cognitively impaired (MCI/mild AD). The AI probability was also converted to a score between 0-100 using the following equation:

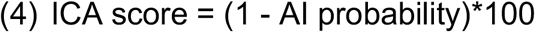

The ICA AI model used in this study was a binary logistic regression machine learning model which is a supervised linear classifier implemented on Python scikit-learn with stochastic gradient descent learning.^55^ The algorithm’s task is to learn a set of weights from a regression model that maps the participant’s ICA test results and demographics to the classification label of healthy or cognitively impaired (Figure 1). An example results-page from the ICA, showing the ICA Score obtained from the AI model, as well as the informative features of the ICA test such as the accuracy, speed and ICA Index are shown in Figure S2 in the SI.

**Figure 1.**
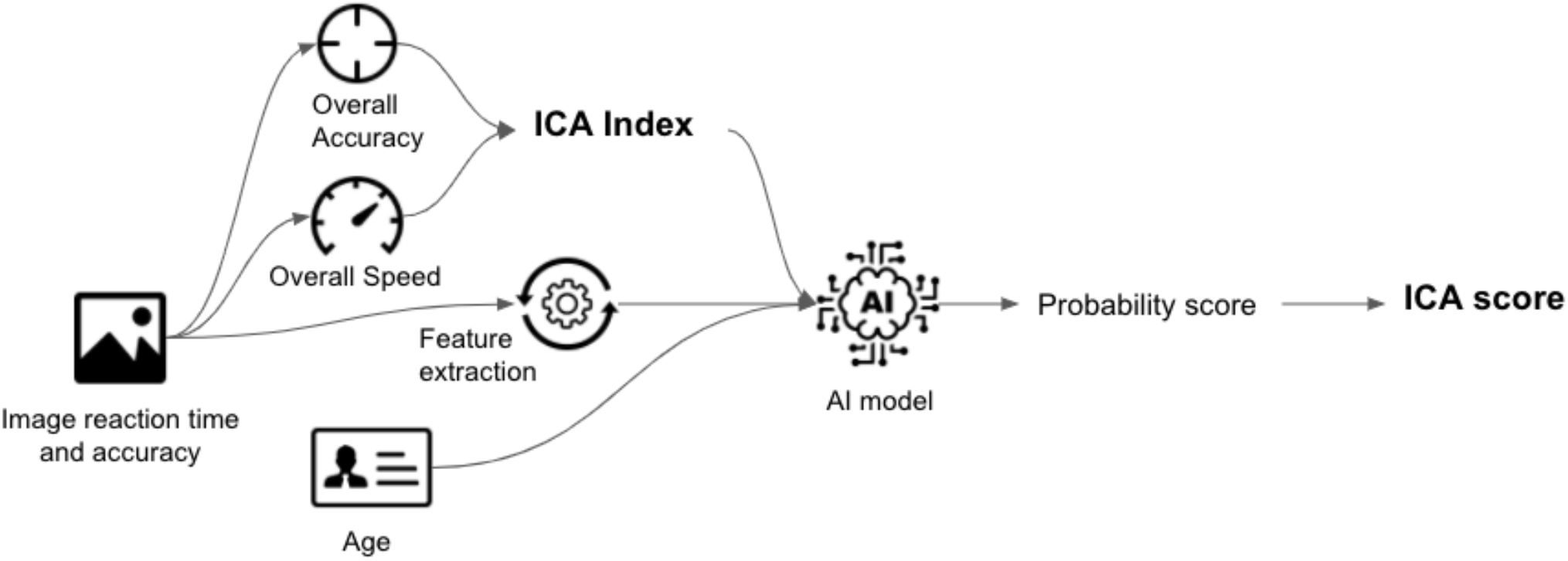
Data features from the ICA test, as well as age are used as features to train the AI model. The trained model is able to give predictions on new unseen data. The AI model outputs a probability score between 0 to 1, which is converted to an ICA Score

The ICA’s prediction on each participant was obtained using leave-one-out-cross validation on the data from Cohort 1 and Cohort 2. In this method the dataset was repeatedly split into two non-overlapping training and testing sets. The training set was used to train the model and the test set to test the performance of the classifier. In the leave-one-out method only one instance was placed in the test set, with the remaining data points used for training. This was done iteratively for all data points, providing an AI probability score for each participant.

The combined results were used to calculate overall metrics (Receiver operating curve area under the curve (ROC AUC), sensitivity, specificity) for the classifier by comparing the ICA AI prediction to clinical diagnosis. The sensitivity or true positive rate is the proportion of actual positives - i.e., impaired-that are identified as such, whilst the specificity, or true negative rate is the proportion of actual negatives - i.e., healthy - that are identified as such.

The ICA AI prediction was also compared to MoCA, ACE to obtain percentage agreement values between these cognitive tests. Single cut-off values were used to obtain predictions for MoCA (score of >=26 for healthy) and ACE (score of >=90 for healthy).

#### 2.4.3. Generalisability of the ICA, and impact of training data size on classification performance

In order to test the generalisability of the ICA, data from Cohort 1 was used to train the AI model and was tested on data from Cohort 2, and vice versa. The number of data points used to train the AI model can significantly impact the performance of the model on the testing dataset. To investigate this, subsets of the data from one cohort were used for training through random sampling. For each training size a model was trained and tested on all the data from the other cohort. We varied the size of the training data from 3 data points to training with all of the data from each cohort.

#### 2.4.4. Assessment of ICA practice effect

To investigate the practice effect related to the ICA, 12 healthy participants (range of 26 – 73, mean 48.2 standard deviation 17.1 years) took 78 tests on a regular basis (936 tests in total). Participants were trained remotely, with assistance provided as needed to initiate the test platform. Thereafter all tests were taken independently at a time and place of the participant’s choosing. Reminders via electronic correspondence were sent periodically to users to encourage adherence.

The time taken for users to complete the 78 tests was 96.8 days on average (standard deviation of 31.7 days). Participants self-administered the ICA remotely, on Apple iPhone devices.

## 3. Results

### 3.1. Baseline characteristics of participants

In total 230 participants (Healthy: 95, MCI: 80, mild AD: 55) were recruited into Cohort 1 and Cohort 2. Participant demographics and cognitive test results are shown in Table 1. Participants were recruited based on a sampling matrix in order to minimise age, gender, and education year differences across the three arms.

Healthy participants did have a lower age compared to mild AD participants. This is reflective of the lower prevalence of young mild AD patients in the general population. There was no significant statistical difference in education years between any of the groups; in all cases two sample t-test p-value was >0.01. For complete table of t-test p-values for age and education years see Table S1 in the SI.

Due to the balanced recruitment, there was no significant difference across genders in any of the cognitive tests (See Figure S3 in SI). The ICA also did not show a significant difference in score between those with 0-11 years education, compared to those with 12 years of education or more (Figure S3b in SI). In contrast there was a statistically significant increase in MoCA score for mild AD participants of higher education, while ACE scores were higher for those with higher education years in Healthy and MCI participants (Table 2).

**Table 2.**
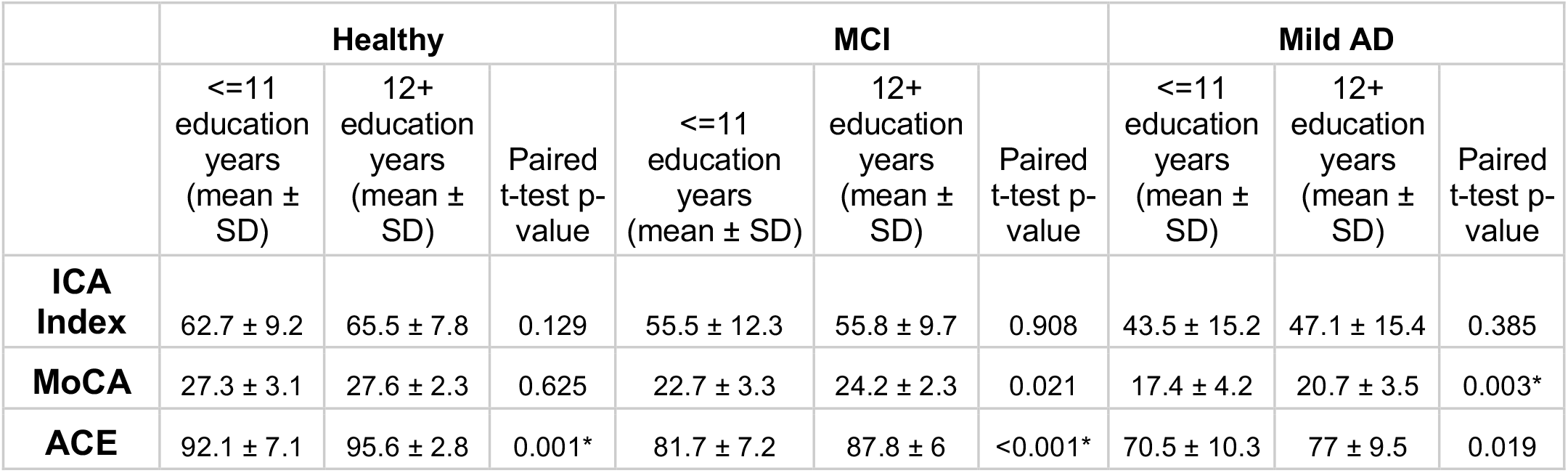
Mean and standard deviation of ICA, MoCA and ACE scores broken down by education years. P-value of t-test between those with 0-11 years education, and those with 12+ years of education, for each cognitive test across the three arms. * indicates significant difference after Bonferroni correction for multiple comparisons

This trend was also illustrated in correlation analysis. The ICA displayed a Pearson r correlation of 0.17 with education years (p=0.01), which is considerably smaller than that of MoCA (r=0.34, p<0.0001) and ACE (r=0.41, p<0.0001) which displayed significant correlations.

### 3.2. ICA convergent validity with MoCA and ACE

The statistically significant Pearson correlation of 0.62 with ACE and 0.58 with MoCA demonstrates convergent validity of ICA with these cognitive tests. The scatterplot of ICA with MoCA and ACE is shown in Figure S4 in the SI. A ceiling effect was observed for MoCA and ACE as a high proportion of healthy participants have maximum test scores, something not observed for the ICA. However, none of the tests were observed to have floor effects, including in the mild AD group.

The breakdown of the ICA Index correlation with the individual cognitive domains as measured by ACE is shown in Table S2 in the SI. In all domains, the Pearson correlation is >0.3, with the strongest correlation obtained with the Memory and Fluency component and a weakest correlation with language.

### 3.3. Speed and accuracy of processing visual information

The breakdown of speed and accuracy by age and diagnosis is shown in Table 3. Within Healthy participants, there is a strong negative correlation between age and accuracy (Pearson r = −0.4, p<0.0001); similarly, for MCI participants (Pearson r = −0.4, p<0.001). However, for mild AD participants there is no correlation between their accuracy and age (Pearson r = 0.07, p=0.58). Analysis did not reveal a significant difference in speed with age within the three groups.

**Table 3.** Mean speed and accuracy on the ICA test, by age category and diagnosis

|  |  | Speed |  | Accuracy |  |
| --- | --- | --- | --- | --- | --- |
|  | Age | mean | std | mean | std |
| Healthy | <70 | 76.3 | 8.4 | 85.9 | 7.6 |
| MCI |  | 73.7 | 11.1 | 78.9 | 9.6 |
| mild AD |  | 70.7 | 18.0 | 64.8 | 17.6 |
| Healthy | $\geq 70$ | 78.3 | 8.6 | 80.6 | 7.6 |
| MCI |  | 71.3 | 14.0 | 74.5 | 11.8 |
| mild AD |  | 69.6 | 16.6 | 65.6 | 15.3 |

Healthy participants have a significantly higher accuracy compared to MCI and mild AD participants in the under 70 age category (t-test p<0.001 after Bonferroni correction for multiple comparisons). MCI participants also had a significantly higher accuracy compared to mild AD in the under 70 age category (t-test p<0.001). In the over 70 age category, healthy participants had significantly higher accuracy compared to mild AD (t-test p<0.001). The other pairwise comparisons by age category and diagnosis were not found to be significant after Bonferroni correction.

Overall, across all ages the Cohen’s D between healthy and MCI participants for accuracy is 0.72 (p<0.0001), and between healthy and mild AD participants it is 1.46 (p<0.0001). Cohen’s D value is 0.41 (p=0.006) for speed (Healthy vs MCI across all ages), compared to 0.52 (p=0.001) for healthy vs mild AD participants. The Cohen’s D for MoCA is 1.43 for Healthy vs MCI (p<0.0001), and 2.37 (p<0.0001) for Healthy vs mild AD. While Cohen’s D measures the effect size, in section 3.4 we present results on comparing the diagnostic accuracy of ICA and >MoCA in relation to clinical diagnosis.

Prior to the commencement of the 100 image ICA test, participants are shown a set of trial images for training purposes. If users perform adequately well on the trial images, they proceed to the main test, however if they perform below chance then the trial images are re-shown to participants. We observed that the number of attempts required by participants before proceeding on to the main test is itself a strong predictor of cognitive impairment. Among Healthy participants 88% completed the trial images on their first attempt compared to 61% of MCI participants, and 44% of mild AD participants (See Figure S5 in SI).

Furthermore, within each group, those who required more than one attempt to progress onto the main test scored lower than those who did not, and they tended to be older participants (Table S3 in SI), indicating within-group cognitive performance variation.

### 3.4. ICA accuracy in detecting cognitive impairment

The raw data from the ICA test consist of categorisation accuracy and reaction time for each of the 100 images on the test. In Figure 2, the average accuracy and reaction time per image has been visualised as a heatmap for each group to show how healthy and impaired participants (MCI and Mild-AD) perform on the ICA test. The sequence of images shown to users during the test is randomised, therefore for ease of comparison the images have been ordered by their category of animal, non-animal.

**Figure 2.**
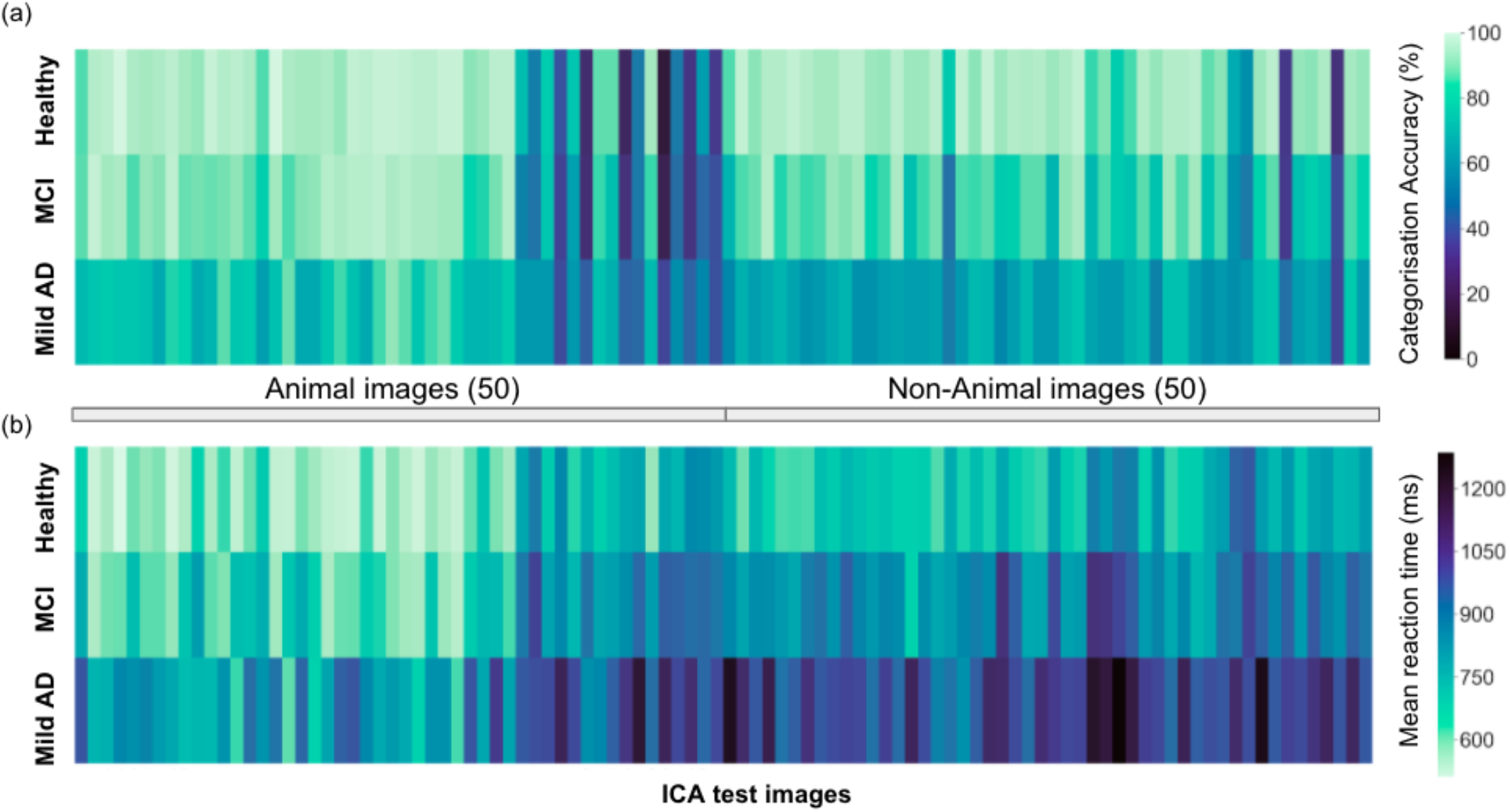
The mean (a) categorisation accuracy (b) reaction time for participants of each diagnosis group, across all the images from the ICA test

Healthy participants display significantly higher categorisation accuracy (Figure 2a) and significantly lower mean reaction time (Figure 2b). However, the varying difficulty of individual images results in a spread of categorisation accuracy for all three groups.

The participants’ age, ICA Index, and features based on the speed and accuracy of responses to the categorisation task extracted from the ICA test were used to train a binomial logistic regression model for classification of healthy vs. MCI/mild AD participants.

Leave one out cross-validation (LOOCV - see methods for full description) was used to obtain a probability of impairment and predictions for each participant. This method ensures the maximum amount of data is used for training the model, while ensuring a separation between the training and test data points. A threshold value of 0.5 was used as the cut-off between healthy and impaired participants.

Figure 3a,b shows the ROC and confusion matrix for distinguishing healthy from impaired (MCI/mild AD) participants. The AUC is 0.84 for distinguishing healthy from impaired with a sensitivity of 79% and specificity of 75%.

**Figure 3.**
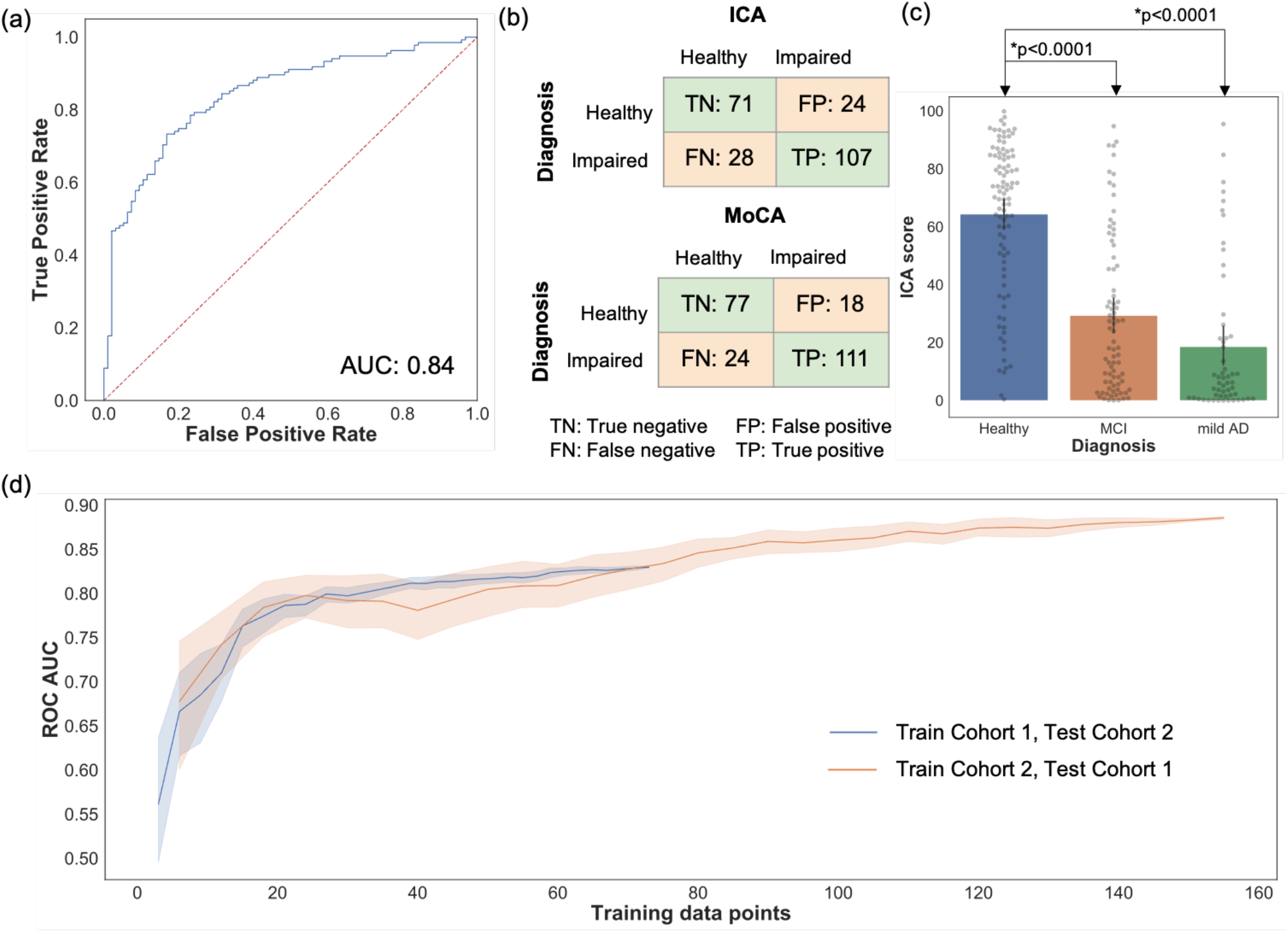
(a) Healthy vs Impaired ROC: AI classification performance by LOOCV (b) The confusion matrix for the ICA and MoCA, comparing the prediction of the cognitive tests with clinical diagnosis. (c) Bar plot with 95% confidence interval of ICA Score for healthy, MCI, and mild AD, with all data points overlaid on the graph. T-test p-value comparing Healthy-MCI, and Healthy - mild AD ICA score is also shown (d) ROC AUC vs training data size. The shaded area represents 95% CI as each training subset was selected randomly 20 times from the whole study data.

For the AI score a higher value is indicative of being cognitively healthy, and a lower score is indicative of potential cognitive impairment. Healthy participants have significantly higher ICA score compared to MCI and mild AD participants (Figure 3c).

A comparison of the classification performance between the ICA and MoCA is shown in Table S4 of the SI. A score of greater than or equal to 26 is the MoCA cut-off for healthy participants, and lower than 26 MoCA cut-off for cognitive impairment, as outlined by Nasreddine et al.^48^ A recent systematic review of pen and paper tests showed that across 20 studies, for healthy versus MCI, MoCA displayed AUC of 85.1%, specificity of 74.6%, and sensitivity of 83.9%.^56^

A direct comparison of this type is not provided for ACE, as this cognitive test was used as an inclusion criterion for healthy participants in Cohort 2, and hence by default it would have a specificity of 100% for healthy participants from that study.

Table S5 in the SI demonstrates the percent agreement between the ICA and MoCA/ACE prediction. Cut off values of greater than or equal to 26 for MoCA, and 90 for ACE, were used to obtain predictions from these tests. In both cases the overall percent agreement is >73%, with the positive percent agreement (where both tests predicted impaired) higher than the negative percent agreement (where both tests predicted healthy). It should be noted that agreement here does not imply correct prediction, as the cognitive tests themselves can misclassify participants.

The ICA AI model has been made explainable by utilising representative, and clinically relevant data from clinical and research studies for training and testing of the model. An inherently more understandable learning algorithm (logistic regression) has been used in favour of more complex ‘black box’ models such as deep learning. An example results page from the ICA is shown in Figure S2 of the SI. In addition to the AI output (ICA Score), the overall accuracy, speed, ICA Index and performance during the test is displayed. As shown in the results presented here, these additional metrics are highly correlated with diagnosis, clinically informative, and help explain the AI output (ICA score), providing supporting evidence to aid the clinician in diagnosis.

### 3.5 ICA AI model generalisability

Figure 3d demonstrates how changing the training sample size impacts the ICA AI model classification accuracy as measured by ROC AUC. Randomly selected subsets of data from Cohort 1 were used as training data, and tested on all of Cohort 2 data, and vice-versa. With small training data sets there is significant fluctuation in performance with wide confidence intervals. Increasing the number of training data points increases the ROC AUC. Furthermore, this analysis demonstrates the generalisability of the ICA AI-model and therefore the ICA test score across two demographically similar populations.

### 3.6 Assessment of ICA practice effect on remote monitoring of cognition

To assess practice effect, we recruited healthy participants to control for the risk of fluctuating or progressively lower test scores in cognitively impaired individuals. The mean ICA Index of the 12 healthy participants, with the 95% confidence interval is shown in Figure S6. The one-way ANOVA p-value obtained was 0.99, showing no significant practice effect for the participants who completed the ICA test 78 times over a period of 96.8 days on average.

## 4. Discussion

In this study we show that the ICA establishes convergent validity with standard-of-care cognitive assessments such as MoCA and ACE. In contrast to these tests the ICA is not confounded by varying levels of education. Similarly, in previous studies conducted in MS patients and healthy controls (174 participants in total) and another study with 436 participants on individuals aged 19 to 98, ICA was shown to have no significant correlation with education years. ^29,30^

The ICA can generalise across populations without the need for collection of population-specific normative data. This was demonstrated by the ability of the ICA to detect patterns of cognitive impairment that are common across cohorts of different cultural and demographic characteristics. Conventional pen and paper and computerised tests require renorming and validation in different languages in order to be validated, requiring collection of culture-specific normative data before a test can be used in populations with different demographic characteristics. Both are prerequisites for large population deployment and risk-based screening in primary care.

We show that the ICA demonstrates no practice effect in healthy participants. As patients with MCI can improve, remain stable, or decline cognitively over time, it is vital that they are monitored regularly for changes in their cognitive status, which could alter diagnosis and management of their care.^3^ MCI monitoring can enable a timely diagnosis and treatment. Available interventions can improve the trajectory of symptoms and the family’s ability to cope with them, and thus change the experience of the course of dementia. ^57^

The conventional method for classification in cognitive assessments is defining a single cut-off value from the test score. This may lead to diagnostic misclassification as single, static cut-offs cannot sufficiently account for factors such as the patient’s age, education level, cognitive reserve and premorbid IQ. The high-dimensional dataset generated by the ICA, as well as the ability to incorporate demographic features, provide more parameters that can be optimised, enabling a classifier to find the optimum classification boundary in higher dimensional space. The ICA’s classification accuracy can be further improved over time by training on additional data to update the AI model.

The ICA utilises an AI model that can be expanded to include and analyse additional patient data such as medication, sleep, and other lifestyle factors along with other biomarker data in order to improve its accuracy and support the development of predictive models of neurodegeneration.

IPS data captured by the ICA show different signature patterns between healthy, MCI and mild AD patients and generate a rich enough dataset to train the ICA AI model to distinguish healthy from cognitively impaired individuals. Subtle changes in IPS can remain undetected, unless rigorously assessed. We are not aware of another cognitive test that quantifies IPS changes to the degree of milliseconds.

The use of AI in decision making, particularly for diagnostic decisions in healthcare, requires a level of explainability from the model which can be used to understand the important factors which led to its output.^58^ This level of explainability can give clinicians confidence in the model, protect against bias and can be used to improve the performance of the system over time. This is in contrast to high accuracy ‘black box’ models that offer limited interpretability of results and therefore prohibit their use in clinical practice.

The ICA results are automatically calculated, eliminating the clinical time required for test interpretation while minimising transcription errors. Test results can be integrated in electronic health records or research databases, an important capability at the intersection between primary and secondary care. The ICA’s ease of use and short duration can improve pre-screening and accelerate participant selection in clinical trials.

Study limitations include a relatively lower recruitment of young participants with mild AD. However, this is reflective of the lower prevalence of young mild AD patients in the general public. Test-retest data have not been captured in this study. We have previously reported that high test-retest reliability (Pearson r > 0.91) was obtained for the ICA.^29,30^

Fluid or molecular biomarker sub-typing to determine amyloid positivity for MCI participants has not been carried in this study, due to lack of data availability. The MCI group, however, reflects the heterogeneity MCI diagnoses in memory clinics. We plan to correlate fluid biomarker positivity with the ICA in future studies.

Remote cognitive assessment is becoming increasingly important, particularly as health services cannot accommodate regular patient attendance to memory services for progression monitoring or response to treatments. The COVID-19 pandemic has accelerated this pressing need and guidelines for the implementation of partly or fully remote memory clinics have recently been published.^16^ Digital cognitive and functional biomarkers are essential in order to enable this. We report a proof-of-concept capability of the ICA for the remote measurement of cognitive performance. Further validation is required for remote administration in MCI and mild AD patients.

In summary the ICA can be used as a digital cognitive biomarker for the detection of MCI and AD. Furthermore, the ICA can be used as a high frequency monitoring tool both in the clinic and potentially remotely. The employment of AI modelling has the potential to further enhance its performance but also to personalise its results at an individual patient level across geographic boundaries.

## Supporting information

All Supplementary Material

## Acknowledgements

We thank the site teams at the NHS trusts and the Royan Institute for their support throughout the study.

## Conflicts

SMKR serves as the Chief Scientific Officer at Cognetivity Ltd. CK serves as the Chief Medical Officer at Cognetivity Ltd and Principal Investigator on NIHR and Industry-funded clinical trials. HM is Data Science Lead at Cognetivity Ltd. DA has received research support and/or honoraria from Astra-Zeneca, H. Lundbeck, Novartis Pharmaceuticals, Biogen, and GE Health, and served as paid consultant for H. Lundbeck, Eisai, Heptares, and Mentis Cura. Other authors declared no potential conflicts of interest.

## References

1. Alzheimer’s Association. 2019 Alzheimer’s disease facts and figures. Alzheimer’s Dement. 2019;15(3):321–387. doi:10.1016/j.jalz.2019.01.010

2. Sachdev PS, Lipnicki DM, Kochan NA, et al. The prevalence of mild cognitive impairment in diverse geographical and ethnocultural regions: The COSMIC Collaboration. PLoS One. 2015;10(11):1–19. doi:10.1371/journal.pone.0142388

3. Petersen RC, Lopez O, Armstrong MJ, et al. Practice guideline update summary: Mild cognitive impairment. Neurology. 2018;90(3):126–135. doi:10.1212/WNL.0000000000004826

4. Alzheimer’s Disease International. World Alzheimer Report 2019.; 2019.

5. Ritchie CW, Russ TC, Banerjee S, et al. The Edinburgh Consensus: preparing for the advent of disease-modifying therapies for Alzheimer’s disease. Alzheimers Res Ther. 2017;9(1):85. doi:10.1186/s13195-017-0312-4

6. Schneider L. A resurrection of aducanumab for Alzheimer’s disease. Lancet Neurol. 2020;19(2):111–112. doi:10.1016/S1474-4422(19)30480-6

7. Dunne RA, Aarsland D, O’Brien JT, et al. Mild cognitive impairment: the Manchester consensus. Age Ageing. Published online November 17, 2020:1–9. doi:10.1093/ageing/afaa228

8. Sabbagh MN, Boada M, Borson S, et al. Rationale for Early Diagnosis of Mild Cognitive Impairment (MCI) supported by Emerging Digital Technologies. J Prev Alzheimer’s Dis. 2020;7(3):1–7. doi:10.14283/jpad.2020.19

9. Sabbagh MN, Boada M, Borson S, et al. Early Detection of Mild Cognitive Impairment (MCI) in an At-Home Setting. J Prev Alzheimer’s Dis. 2020;7(3):171–178. doi:10.14283/jpad.2020.22

10. Duboisa B, Padovanib A, Scheltensc P, Rossid A, Agnello GD. Timely diagnosis for alzheimer’s disease: A literature review on benefits and challenges. J Alzheimer’s Dis. 2015;49(3):617–631. doi:10.3233/JAD-150692

11. Clare L, Wu YT, Teale JC, et al. Potentially modifiable lifestyle factors, cognitive reserve, and cognitive function in later life: A cross-sectional study. PLoS Med. 2017;14(3):1–14. doi:10.1371/journal.pmed.1002259

12. Xu G, Meyer JS, Thornby J, Chowdhury M, Quach M. Screening for mild cognitive impairment (MCI) utilizing combined mini-mental-cognitive capacity examinations for identifying dementia prodromes. Int J Geriatr Psychiatry. 2002;17(11):1027–1033. doi:10.1002/gps.744

13. de Jager CA, Schrijnemaekers ACMC, Honey TEM, Budge MM. Detection of MCI in the clinic: Evaluation of the sensitivity and specificity of a computerised test battery, the Hopkins Verbal Learning Test and the MMSE. Age Ageing. 2009;38(4):455–460. doi:10.1093/ageing/afp068

14. Ranson JM, Kuźma E, Hamilton W, Muniz-Terrera G, Langa KM, Llewellyn DJ. Predictors of dementia misclassification when using brief cognitive assessments. Neurol Clin Pract. 2019;9(2):109–117. doi:10.1212/CPJ.0000000000000566

15. Cooley SA, Heaps JM, Bolzenius JD, et al. Longitudinal Change in Performance on the Montreal Cognitive Assessment in Older Adults. Clin Neuropsychol. 2015;29(6):824–835. doi:10.1080/13854046.2015.1087596

16. Owens AP, Ballard C, Beigi M, et al. Implementing Remote Memory Clinics to Enhance Clinical Care During and After COVID-19. Front Psychiatry. 2020;11(September):1–15. doi:10.3389/fpsyt.2020.579934

17. Zygouris S, Giakoumis D, Votis K, et al. Can a virtual reality cognitive training application fulfill a dual role? Using the virtual supermarket cognitive training application as a screening tool for mild cognitive impairment. J Alzheimer’s Dis. 2015;44(4):1333–1347. doi:10.3233/JAD-141260

18. Hayes TL, Abendroth F, Adami A, Pavel M, Zitzelberger TA, Kaye JA. Unobtrusive assessment of activity patterns associated with mild cognitive impairment. Alzheimer’s Dement. 2008;4(6):395–405. doi:10.1016/j.jalz.2008.07.004

19. Weintraub S, Dikmen SS, Heaton RK, et al. The Cognition Battery of the NIH Toolbox for Assessment of Neurological and Behavioral Function: Validation in an Adult Sample. J Int Neuropsychol Soc. 2014;20(6):567–578. doi:10.1017/S1355617714000320

20. Erlanger DM, Kaushik T, Broshek D, Freeman J, Feldman D, Festa J. Development and validation of a web-based screening tool for monitoring cognitive status. J Head Trauma Rehabil. 2002;17(5):458–476. doi:10.1097/00001199-200210000-00007

21. Kourtis LC, Regele OB, Wright JM, Jones GB. Digital biomarkers for Alzheimer’s disease: the mobile/wearable devices opportunity. npj Digit Med. 2019;2(1):9. doi:10.1038/s41746-019-0084-2

22. Jo T, Nho K, Saykin AJ. Deep Learning in Alzheimer’s Disease: Diagnostic Classification and Prognostic Prediction Using Neuroimaging Data. Front Aging Neurosci. 2019;11(August). doi:10.3389/fnagi.2019.00220

23. Park JH, Cho HE, Kim JH, et al. Machine learning prediction of incidence of Alzheimer’s disease using large-scale administrative health data. npj Digit Med. 2020;3(1). doi:10.1038/s41746-020-0256-0

24. Tanveer M, Richhariya B, Khan RU, et al. Machine learning techniques for the diagnosis of alzheimer’s disease: A review. ACM Trans Multimed Comput Commun Appl. 2020;16(1s). doi:10.1145/3344998

25. Alzheimer’s Research UK. Dementia Attitudes Monitor.; 2018. www.dementiastatistics.org/attitudes

26. Parsey CM, Schmitter-Edgecombe M. Applications of Technology in Neuropsychological Assessment. Clin Neuropsychol. 2013;27(8):1328–1361. doi:10.1080/13854046.2013.834971

27. Koo BM, Vizer LM. Mobile Technology for Cognitive Assessment of Older Adults: A Scoping Review. Innov Aging. 2019;3(1):1–14. doi:10.1093/geroni/igy038

28. Fazio S, Pace D, Flinner J, Kallmyer B. The Fundamentals of Person-Centered Care for Individuals with Dementia. Gerontologist. 2018;58:S10–S19. doi:10.1093/geront/gnx122

29. Khaligh-Razavi S-M, Habibi S, Sadeghi M, et al. Integrated Cognitive Assessment: Speed and Accuracy of Visual Processing as a Reliable Proxy to Cognitive Performance. Sci Rep. 2019;9(1):1102. doi:10.1038/s41598-018-37709-x

30. Khaligh-Razavi S-M, Sadeghi M, Khanbagi M, Kalafatis C, Nabavi SM. A self-administered, artificial intelligence (AI) platform for cognitive assessment in multiple sclerosis (MS). BMC Neurol. 2020;20(1):193. doi:10.1186/s12883-020-01736-x

31. Kiani R, Esteky H, Mirpour K, Tanaka K. Object Category Structure in Response Patterns of Neuronal Population in Monkey Inferior Temporal Cortex. J Neurophysiol. 2007;97(6):4296–4309. doi:10.1152/jn.00024.2007

32. Cichy RM, Pantazis D, Oliva A. Resolving human object recognition in space and time. Nat Neurosci. 2014;17(3):455–462. doi:10.1038/nn.3635

33. Karimi H, Marefat H, Khanbagi M, Kalafatis C. Temporal dynamics of animacy categorization in the brain of patients with mild cognitive impairment. bioRxiv. Published online 2020:0-20. doi:https://doi.org/10.1101/2020.11.20.390435

34. Kriegeskorte N, Mur M, Ruff DA, et al. Matching Categorical Object Representations in Inferior Temporal Cortex of Man and Monkey. Neuron. 2008;60(6):1126–1141. doi:10.1016/j.neuron.2008.10.043

35. Naselaris T, Stansbury DE, Gallant JL. Cortical representation of animate and inanimate objects in complex natural scenes. J Physiol. 2012;106(5-6):239–249. doi:10.1016/j.jphysparis.2012.02.001

36. Liu H, Agam Y, Madsen JR, Kreiman G. Timing, Timing, Timing: Fast Decoding of Object Information from Intracranial Field Potentials in Human Visual Cortex. Neuron. 2009;62(2):281–290. doi:10.1016/j.neuron.2009.02.025

37. Khaligh-Razavi S-M, Cichy RM, Pantazis D, Oliva A. Tracking the Spatiotemporal Neural Dynamics of Real-world Object Size and Animacy in the Human Brain. J Cogn Neurosci. 2018;30(11):1559–1576. doi:10.1162/jocn_a_01290

38. Fahrenfort JJ, Scholte HS, Lamme VAF. Masking Disrupts Reentrant Processing in Human Visual Cortex. J Cogn Neurosci. 2007;19(9):1488–1497. doi:10.1162/jocn.2007.19.9.1488

39. Rajaei K, Mohsenzadeh Y, Ebrahimpour R, Khaligh-Razavi S-M. Beyond core object recognition: Recurrent processes account for object recognition under occlusion. Isik L, ed. PLOS Comput Biol. 2019;15(5):e1007001. doi:10.1371/journal.pcbi.1007001

40. Mirzaei A, Khaligh-Razavi S-M, Ghodrati M, Zabbah S, Ebrahimpour R. Predicting the human reaction time based on natural image statistics in a rapid categorization task. Vision Res. 2013;81:36–44. doi:10.1016/j.visres.2013.02.003

41. Marx S, Hansen-Goos O, Thrun M, Einhauser W. Rapid serial processing of natural scenes: Color modulates detection but neither recognition nor the attentional blink. J Vis. 2014;14(14):4–4. doi:10.1167/14.14.4

42. Zhu W, Drewes J, Gegenfurtner KR. Animal Detection in Natural Images: Effects of Color and Image Database. de Beeck HPO, ed. PLoS One. 2013;8(10):e75816. doi:10.1371/journal.pone.0075816

43. Bacon-Macé N, Macé MJ-M, Fabre-Thorpe M, Thorpe SJ. The time course of visual processing: Backward masking and natural scene categorisation. Vision Res. 2005;45(11):1459–1469. doi:10.1016/j.visres.2005.01.004

44. Khaligh-Razavi S-M, Kriegeskorte N. Deep Supervised, but Not Unsupervised, Models May Explain IT Cortical Representation. Diedrichsen J, ed. PLoS Comput Biol. 2014;10(11):e1003915. doi:10.1371/journal.pcbi.1003915

45. Costa SL, Genova HM, DeLuca J, Chiaravalloti ND. Information processing speed in multiple sclerosis: Past, present, and future. Mult Scler J. 2017;23(6):772–789. doi:10.1177/1352458516645869

46. DeLuca J, Chelune GJ, Tulsky DS, Lengenfelder J, Chiaravalloti ND. Is Speed of Processing or Working Memory the Primary Information Processing Deficit in Multiple Sclerosis? J Clin Exp Neuropsychol. 2004;26(4):550–562. doi:10.1080/13803390490496641

47. Lu H, Chan SSM, Lam LCW. ‘Two-level’ measurements of processing speed as cognitive markers in the differential diagnosis of DSM-5 mild neurocognitive disorders (NCD). Sci Rep. 2017;7(1):521. doi:10.1038/s41598-017-00624-8

48. Nasreddine ZS, Phillips NA, Bédirian V, et al. The Montreal Cognitive Assessment, MoCA: a brief screening tool for mild cognitive impairment. J Am Geriatr Soc. 2005;53(4):695–699. doi:10.1111/j.1532-5415.2005.53221.x

49. Mathuranath PS, Nestor PJ, Berrios GE, Rakowicz W, Hodges JR. A brief cognitive test battery to differentiate Alzheimer’s disease and frontotemporal dementia. Neurology. 2000;55(11):1613–1620. doi:10.1212/01.wnl.0000434309.85312.19

50. Hodges JR, Larner AJ. Addenbrooke’s Cognitive Examinations: ACE, ACE-R, ACE-III, ACEapp, and M-ACE. In: Cognitive Screening Instruments. Springer International Publishing; 2017:109–137. doi:10.1007/978-3-319-44775-9_6

51. Mioshi E, Dawson K, Mitchell J, Arnold R, Hodges JR. The Addenbrooke’s Cognitive Examination Revised (ACE-R): a brief cognitive test battery for dementia screening. Int J Geriatr Psychiatry. 2006;21(11):1078–1085. doi:10.1002/gps.1610

52. Hsieh S, Schubert S, Hoon C, Mioshi E, Hodges JR. Validation of the Addenbrooke’s Cognitive Examination III in Frontotemporal Dementia and Alzheimer’s Disease. Dement Geriatr Cogn Disord. 2013;36(3-4):242–250. doi:10.1159/000351671

53. McKhann GM, Knopman DS, Chertkow H, et al. The diagnosis of dementia due to Alzheimer’s disease: Recommendations from the National Institute on Aging-Alzheimer’s Association workgroups on diagnostic guidelines for Alzheimer’s disease. Alzheimer’s Dement. 2011;7(3):263–269. doi:10.1016/j.jalz.2011.03.005

54. Willis BH. Spectrum bias - Why clinicians need to be cautious when applying diagnostic test studies. Fam Pract. 2008;25(5):390–396. doi:10.1093/fampra/cmn051

55. SGDClassifier. scikit-learn. Accessed February 19, 2021. https://scikit-learn.org/stable/modules/generated/sklearn.linear_model.SGDClassifier.html

56. De Roeck EE, De Deyn PP, Dierckx E, Engelborghs S. Brief cognitive screening instruments for early detection of Alzheimer’s disease: A systematic review. Alzheimer’s Res Ther. 2019;11(1):1–14. doi:10.1186/s13195-019-0474-3

57. Livingston G, Sommerlad A, Orgeta V, et al. Dementia prevention, intervention, and care. Lancet. 2017;390(10113):2673–2734. doi:10.1016/S0140-6736(17)31363-6

58. The Royal Society. Explainable AI: The Basics.; 2019. https://royalsociety.org/topics-policy/projects/explainable-ai/

