## Supplementary material for "Validity and cultural generalisability of a 5-minute AI-based, computerised cognitive assessment in Mild Cognitive Impairment and Alzheimer’s Dementia": All Supplementary Material

### Supplementary Information

<u>Cohort 1&2 Eligibility Criteria</u>

**Inclusion Criteria for Healthy group**

1. Bristol Activities of Daily Living Scale (BADLS) score of “unimpaired” or “mildly impaired”
2. Capacity to understand the information about the study and to give consent to participate
3. Males and females aged between 55-90 years.
4. Not currently on medication that may interfere with the study results.
5. In good general health.

Additional inclusion criteria which was only used in Cohort 2:
Addenbrooke’s Cognitive Assessment (ACE-III) score of >= 90/100

**Exclusion Criteria for Healthy group**
1. Presence of significant cerebrovascular disease i.e. history of CVA
2. Major medical comorbidities e.g. Congestive Cardiac Failure, Diabetes Mellitus with renal impairment stage 4 or 5
3. Major psychiatric disorder e.g. chronic psychosis, recurrent depressive disorder, generalised anxiety disorder .
4. The use of cognitive enhancing drugs e.g. cholinesterase inhibitors
5. A concurrent diagnosis of epilepsy
6. A history of alcohol dependence
7. A history of illicit drug use
8. A history of severe visual impairment, e.g. macular degeneration, diabetic retinopathy, as determined by the clinical team
9. Presence of Sleep Apnoea
10. A history of head trauma
11. Severe upper limb arthropathy

Inclusion Criteria for MCI group
1. A clinical diagnosis of MCI according to validated criteria (within the past 12 months, earlier diagnosis of MCI with memory clinic conformation within the past 12 months is also acceptable)
2. Males and females aged 55-90 years.
3. Willing and able to provide informed consent

**Exclusion Criteria for MCI group**
1. Patients who fulfill criteria for a diagnosis of mild AD
2. Major medical comorbidities i.e. Congestive Cardiac Failure, Diabetes Mellitus with renal impairment stage 4 or 5.
3. Major psychiatric disorder e.g. chronic psychosis, recurrent depressive disorder, generalised anxiety disorder.
4. The use of cognitive enhancing drugs e.g. cholinesterase inhibitors

5. A concurrent diagnosis of epilepsy
6. A history of alcohol misuse
7. A history of illicit drug use
8. A history of severe visual impairment, e.g. macular degeneration, diabetic retinopathy, as determined by the clinical team
9. A history of head trauma
10. Presence of Sleep Apnoea
11. Severe upper limb arthropathy
12. Diagnosis of MCI related to a cerebrovascular event (commonly referred to as Vascular Cognitive Impairment)

**Inclusion Criteria for mild AD group**

1. A clinical diagnosis of mild AD according to validated criteria (within the past 12 months, earlier diagnosis of mild AD with memory clinic conformation within the past 12 months is also acceptable)
2. Males and females aged 55-90 years.
3. Willing and able to provide informed consent

**Exclusion Criteria for mild AD group**

1. Patients who fulfill criteria for a diagnosis of Moderate AD or other types of Dementia such as Posterior Cortical Atrophy
2. Major medical comorbidities e.g. Congestive Cardiac Failure, Diabetes Mellitus with renal impairment stage 4 or 5
3. Major psychiatric disorder eg. chronic psychosis, recurrent depressive disorder, generalised anxiety disorder.
4. A concurrent diagnosis of epilepsy
5. A history of alcohol misuse
6. A history of illicit drug use
7. A history of severe visual impairment, e.g. macular degeneration, diabetic retinopathy, as determined by the clinical team
8. A history of head trauma
9. Presence of Sleep Apnoea.
10. Severe upper limb arthropathy

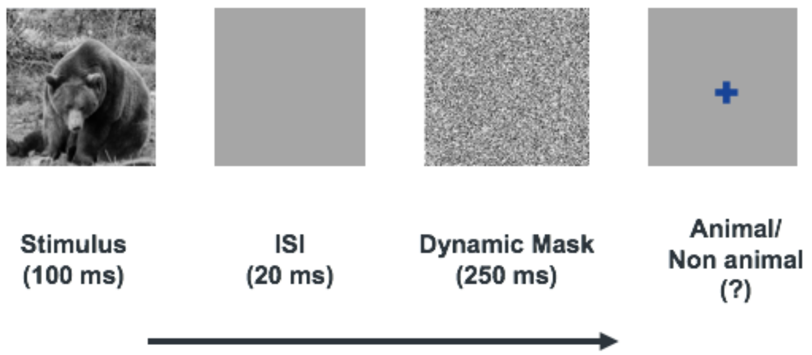


Figure S1. One hundred natural images (50 animal and 50 non-animal) with various levels of difficulty are presented to the participants. Each image is presented for 100 ms followed by 20 ms inter-stimulus interval (ISI), followed by a dynamic noise mask (for 250 ms), followed by subject's categorization into animal vs. non-animal.

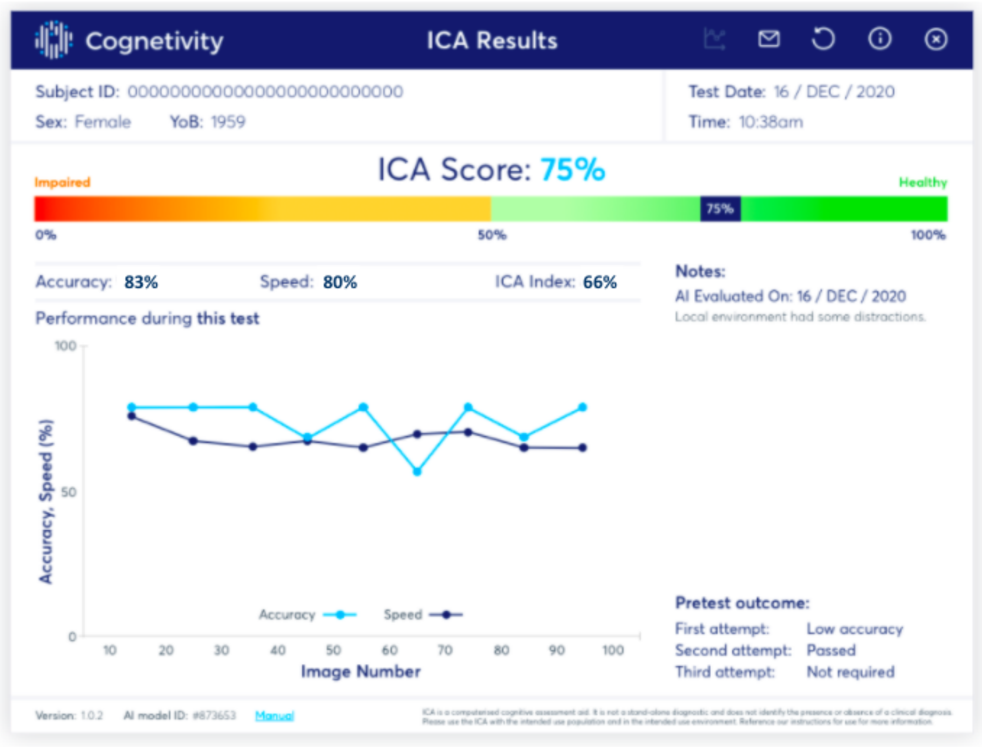


Figure S2. An example results page from the ICA. As well as the ICA Score, breakdown of performance on the test is shown, such as the overall accuracy, speed and ICA Index. The line chart displays how the accuracy and speed of the participants varied during the 100-image test. The pre-tests outcome indicates the performance of the participants on trial images shown before the main test

Table S1. T-test p-value for age and education years, comparing the three arms in the combined dataset. * to indicate a p-value below 0.01.

| **Comparison** | **t-test p-value Age** | **t-test p-value Education years** |
|---|---:|---:|
| Healthy-MCI | 0.018 | 0.067 |
| Healthy-mild AD | 0.001 | 0.039 |
| MCI-mild AD | 0.254 | 0.701 |

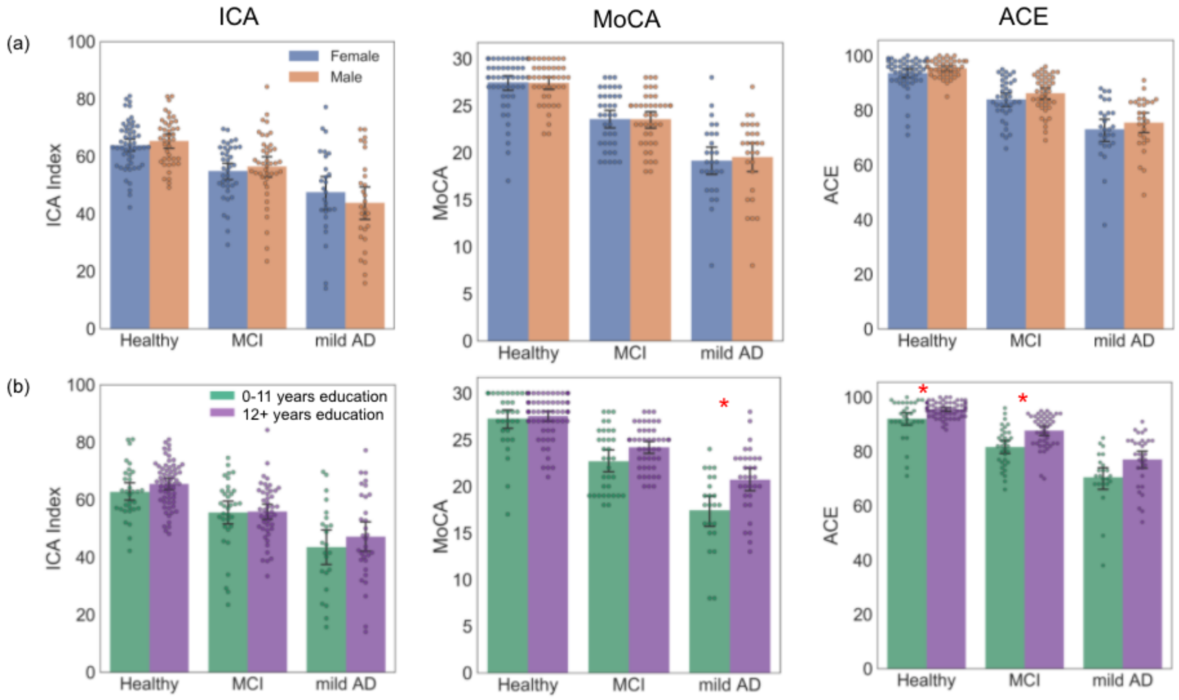


Figure S3. Plots showing the impact of demographic features on ICA Index, MoCA and ACE (a) Gender (b) Education years. Each dot represents one participant. A red star has been used to indicate where significant difference was observed

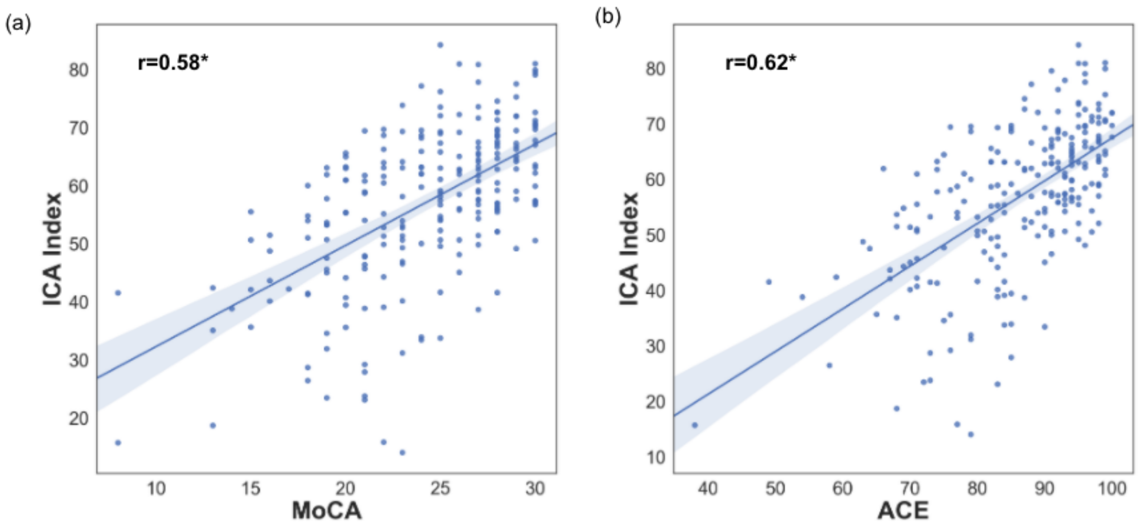


Figure S4. ICA Index correlation with ACE and MoCA (a) Pearson correlation: 0.58, p<0.0001, ICA Score Pearson correlation with MoCA is 0.58, p<0.0001 (b) Pearson correlation 0.62, p<0.0001; ICA Score Pearson correlation with ACE is 0.56, p<0.0001. For breakdown of correlation with ACE subdomains see Table S2 in the SI.

Table S2 Correlation of the ICA with cognitive domains of ACE

| ACE domain | Pearson correlation | Pearson p-value |
|---|---|---|
| Memory | 0.53 | p<0.0001 |
| Attention | 0.41 | p<0.0001 |
| Fluency | 0.53 | p<0.0001 |
| Language | 0.31 | p<0.0001 |
| Visuospatial | 0.48 | p<0.0001 |

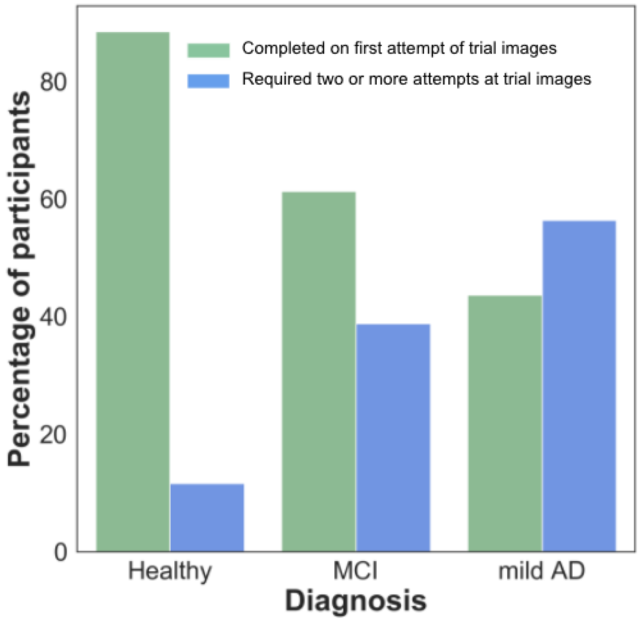


Figure S5. Percentage of participants who completed the main ICA test after successfully completing the pre-test trial images on their first attempt, compared to those who needed two or more attempts at the trial images.

Table S3. Mean ICA Index and Age when one trial attempt was needed, compared to two or more, broken down by diagnosis

| | | | ICA Index | | Age | |
|---|---|---|---|---|---|---|
| Diagnosis | Number of trial attempts | count | mean | std | mean | std |
| Healthy | One | 84 | 65.3 | 8.1 | 66.5 | 7.4 |
| | Two or more | 11 | 58.6 | 8.3 | 69.1 | 9.5 |
| MCI | One | 49 | 60.2 | 8.4 | 67.3 | 6.8 |
| | Two or more | 31 | 48.6 | 10.4 | 73.3 | 8.5 |
| mild AD | One | 24 | 53.6 | 14.0 | 70.5 | 7.3 |
| | Two or more | 31 | 39.4 | 13.3 | 71.8 | 8.4 |

Table S4. Classification performance metrics for ICA and MoCA, with 95% confidence intervals (CI)

| **Cognitive test** | **Classification** | **AUC (95% CI)** | **Sensitivity (95% CI)** | **Specificity (95% CI)** |
|---|---|---|---|---|
| ICA | Healthy vs Impaired | 0.842 (0.791, 0.893) | 79.3 (72.4, 86.1) | 74.7 (66.0, 83.5) |
| ICA | Healthy vs MCI | 0.814 (0.749, 0.878) | 76.2 (66.9, 85.6) | 74.7 (66.0, 83.5) |
| ICA | Healthy vs mild AD | 0.883 (0.823, 0.944) | 83.6 (73.9, 93.4) | 74.7 (66.0, 83.5) |
| MoCA | Healthy vs Impaired | 0.816 (0.765, 0.868) | 82.2 (75.8, 88.7) | 81.1 (73.2, 88.9) |
| MoCA | Healthy vs MCI | 0.768 (0.705, 0.831) | 72.5 (62.7, 82.3) | 81.1 (73.2, 88.9) |
| MoCA | Healthy vs mild AD | 0.887 (0.84, 0.934) | 96.4 (91.4, 100.0) | 81.1 (73.2, 88.9) |

Table S5: Percent agreement between ICA and MoCA, ACE, with 95% confidence intervals

| | **Positive percent agreement (95% CI)** | **Negative percent agreement (95% CI)** | **Overall percent agreement (95% CI)** |
|---|---|---|---|
| **ICA and MoCA prediction** | 77.5 (70.3, 84.7) | 69.3 (60.3, 78.3) | 73.9 (68.2, 79.6) |
| **ICA and ACE prediction** | 81.4 (74.2, 88.6) | 66.7 (58.1, 75.2) | 73.9 (68.2, 79.6) |

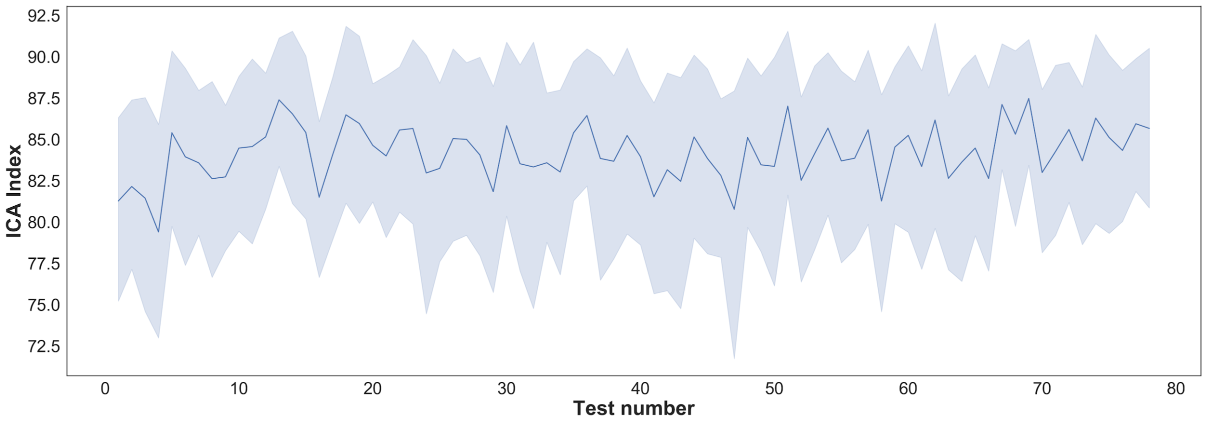


Figure S6. Mean ICA Index of 12 participants over 78 tests, taken on an approximately daily basis. The shaded areas represent 95% CI. ANOVA one way p-value is 0.99
